# Dense neuronal reconstruction through X-ray holographic nano-tomography

**DOI:** 10.1101/653188

**Authors:** Alexandra Pacureanu, Jasper Maniates-Selvin, Aaron T. Kuan, Logan A. Thomas, Chiao-Lin Chen, Peter Cloetens, Wei-Chung Allen Lee

## Abstract

Elucidating the structure of neuronal networks provides a foundation for understanding how the nervous system processes information to generate behavior. Despite technological breakthroughs in visible light and electron microscopy, imaging dense nanometer-scale neuronal structures over millimeter-scale tissue volumes remains a challenge. Here, we demonstrate that X-ray holographic nano-tomography is capable of imaging large tissue volumes with sufficient resolution to disentangle dense neuronal circuitry in *Drosophila melanogaster* and mammalian central and peripheral nervous tissue. Furthermore, we show that automatic segmentation using convolutional neural networks enables rapid extraction of neuronal morphologies from these volumetric datasets. The technique we present allows rapid data collection and analysis of multiple specimens, and can be used correlatively with light microscopy and electron microscopy on the same samples. Thus, X-ray holographic nano-tomography provides a new avenue for discoveries in neuroscience and life sciences in general.

## Introduction

Our understanding of the nervous system is built on our knowledge of the structure and connectivity of neurons. However, unraveling their structure is a daunting technical challenge because neuronal axons and dendrites are small in diameter (~20-1000 nm) and traverse long distances (millimeters or more). With current technology, we generally can either image a sparse subset of neurons with a large field of view (FOV) through fluorescent labeling and visible light microscopy (LM) or comprehensively map small regions via electron microscopy (EM). Advancing our understanding of neuronal networks will require imaging modalities that can simultaneously achieve high resolution and large FOV.

At present, LM and magnetic resonance imaging techniques can image large regions, but not with sufficient resolution to densely resolve neuronal processes. Recent advances in super-resolution imaging have greatly improved resolving power for LM (Gao et al., 2019; Heintzmann and Gustafsson, 2009; Hell, 2007; Huang et al., 2009); however, these techniques rely on sparse labeling with fluorescent probes, such that only a fraction of neurons are visible. On the other hand, all neurons can be detected locally in EM micrographs, but collection of large volumes encompassing entire circuits requires elaborate sample and data processing, yielding major scaling challenges (Briggman and Bock, 2012; Helmstaedter et al., 2013; Lee et al., 2016; Xu et al., 2017; Zheng et al., 2018). Thus, despite rapid and transformative advances in both LM and EM, a spatial resolution and FOV gap remains.

Hard X-rays are an attractive illumination probe for imaging thick tissue samples with fine spatial resolution due to their high penetration power and sub-nanometer wavelength. Previously, attenuation-based X-ray micro-tomography with a synchrotron source enabled visualization of metal stained, sparse wiring in the *Drosophila* brain (Mizutani et al., 2013). The advent of X-ray phase-contrast techniques enabled imaging of both stained and unstained brain tissue at micrometer and sub-micrometer scales (Cedola et al., 2017; Dyer et al., 2017; Fonseca et al., 2018; Khimchenko et al., 2018; Massimi et al., 2019; Schulz et al., 2010; Shahmoradian et al., 2017; Töpperwien et al., 2018). However, until now, X-ray microscopy has not achieved the necessary resolving power to enable dense reconstruction of neuronal morphologies.

Here we present a pipeline based on X-ray holographic nano-tomography (also called X-ray nanoholotomography, or XNH) that enables imaging of large volumes of neural tissue with sufficient resolution to extract the morphologies of densely packed neurons without specific labeling. We combined a highly focused and brilliant hard X-ray nano-probe (da Silva et al., 2017), an advanced nano-positioning system, cryogenic imaging, new sample preparation protocols, and enhanced phase retrieval approaches to achieve significant improvements in image quality. We show that individual neurons can be manually traced through XNH datasets of central and peripheral nervous tissue of the fruit fly *Drosophila melanogaster* and mouse. In addition, we use a 3D U-NET convolutional neural network (CNN) to automatically segment dendrites and axons from the XNH data. Moreover, we demonstrate that XNH is compatible with cell type-specific genetic labeling techniques as well as post-hoc EM of the same samples. These results demonstrate that XNH is a versatile volumetric imaging modality for biological microscopy, which can be combined with LM and EM in correlative workflows to accelerate our understanding of neuronal networks.

## Results

### XNH imaging of central and peripheral nervous systems

To explore the potential of XNH for imaging neuronal morphologies, we imaged samples of mouse cortex as well as the brain, ventral nerve cord (VNC), and leg of adult *Drosophila*. We varied the reconstructed isotropic voxel size (edge length) from 30 nm to 120 nm, yielding FOVs (edge length of square cylindrical volume) from 60 µm to 240 µm (96 µm to 384 µm with extended FOV, see Supplementary Table 1) for each dataset. Sample sizes ranged from whole *Drosophila* brains (300 µm × 200 µm × 700 µm) to blocks of mouse cortex (1 mm × 0.5 mm × 2 mm). Samples were prepared using standard protocols for EM (Hua et al., 2015; Zheng et al., 2018) and mounted onto aluminum cylindrical pins for imaging (Fig. 1a, inset).

**Figure 1:**
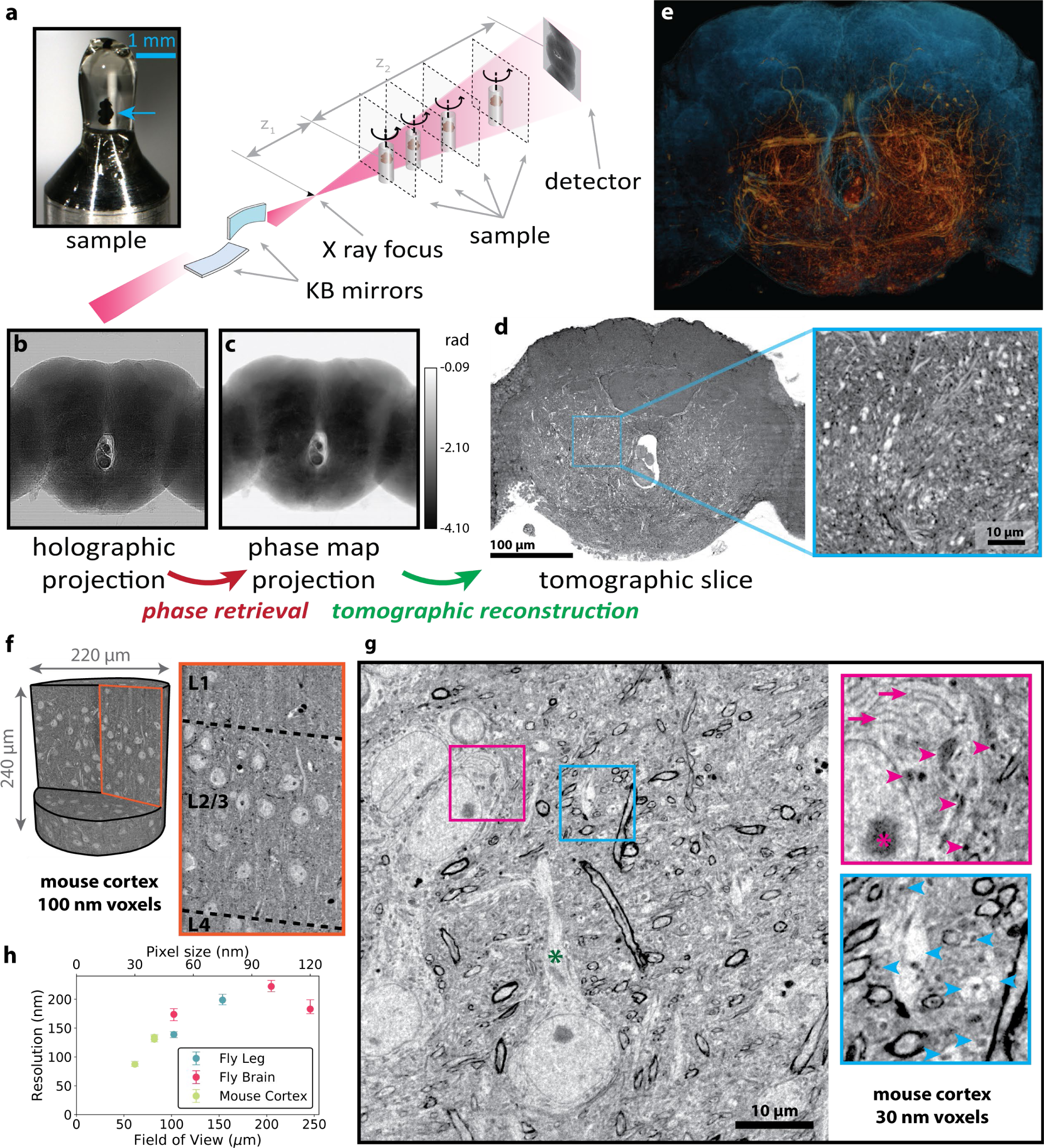
X-ray Holographic Nano-Tomography (XNH) Technique and Characterization. **(a)** Upper inset: a *Drosophila* brain (blue arrow) embedded in resin and mounted on a pin for imaging. Diagram: schematic of imaging setup. The X-ray beam from the synchrotron is focused to a spot using two mirrors, and traverses the sample before hitting the detector. **(b)** Holographic projections (a result of free-space propagation of the coherent X-ray beam) are recorded for each angle as the sample is rotated up to 180°. **(c)** Phase projections are calculated using phase retrieval algorithms by computationally combining 4 holographic projections of the sample at different distances from the beam focus. Color scale indicates phase in radians. **(d)** A slice through the reconstructed 3D image volume (120 nm voxels), calculated using tomographic reconstruction. Inset right: detail view, showing individual neurons (bright objects). **(e)** 3D rendering of XNH volume of the central fly brain. The tissue outline is shown in blue, while large neurons and trachea are highlighted in orange. **(f)** 3D rendering of an XNH volume of mouse cortex (100 nm voxel size). Inset right: virtual slice resolving individual cell bodies and large dendrites. **(g)** Virtual slice through a mouse cortex XNH volume with 30 nm voxels. Insets: detail view resolving ultrastructural features including mitochondria (magenta arrowheads), endoplasmic reticulum (magenta arrows), nucleolus (magenta asterisk), myelinated axons, and dendrites (blue arrowheads). **(h)** Measured resolution for different scans plotted as a function of pixel size and field of view (FOV). Values for resolution were obtained using Fourier Shell Correlation (see Methods, Supplementary Table 1).

XNH imaging was conducted at the nano-imaging beamline ID16A at the European Synchrotron (ESRF). For image acquisition, the samples were positioned downstream of the focal spot of a highly brilliant X-ray beam (Fig. 1a). After traversing the sample, the propagating beam generated self-interference patterns, (i.e. holograms) that were recorded on a lens-coupled CCD detector about 1.2 m downstream of the sample (Fig. 1b). For each dataset, four tomographic scans (rotations of the sample over 180°) were recorded at different focus-to-sample distances (Fig. 1a), and processed together via a phase retrieval algorithm to obtain angular phase maps of the sample (Fig. 1c) (Cloetens et al., 1999; Mokso et al., 2007; Yu et al., 2018). Lastly, a 3D image volume of the tissue was generated from the angular phase maps by tomographic reconstruction using filtered back-projection (Mirone et al., 2014).

Figure 1d-e shows a tomographic slice and a 3D rendering of an XNH scan capturing the central brain of an adult *Drosophila* (120 nm voxels) (Supplementary Video 1). At this resolution, large individual neuronal processes can be resolved (Fig. 1d, inset, Fig. 1e, orange tracts). Figure 1f shows a volume rendering of a large FOV XNH scan from mouse cortex (100 nm voxels). In this dataset, we can trace large dendrites, in particular the apical dendrites of pyramidal neurons. Figure 1g shows a virtual slice from a high-resolution mouse cortex scan (30 nm voxels) (Supplementary Video 2). At this resolution, many ultrastructural features are resolved including mitochondria, endoplasmic reticulum, dendrites, myelinated and unmyelinated axons (Fig. 1g, insets).

To quantify the spatial resolution of XNH image volumes, we utilized Fourier Shell Correlation (FSC) (Harauz and van Heel, 1986). While the pixel size is directly determined by the geometrical magnification M = (*z*_1_ + *z*_2_)/*z*_1_ (Fig. 1a), the actual resolution depends on multiple factors, including the spot size and coherence properties, the mechanical stability of the system, the detection system, the sample composition and the image reconstruction approach. We found that the measured spatial resolutions in both mouse and *Drosophila* data vary between two and four times the voxel size, depending on the sample characteristics and acquisition parameters (Fig. 1h, Supplementary Table 1, Fig. S1a-c). Reducing the voxel size improved image resolution down to the smallest voxel size we tested (30 nm), but the relative resolution (measured resolution divided by voxel size) was best at larger voxel sizes (Fig. S1a). This is likely due to larger voxel sizes corresponding to reduced radiation dose and less tissue present outside the FOV (see Discussion).

Although FSC is a commonly utilized method to quantify resolution in many imaging modalities including X-ray imaging, its implementation is somewhat controversial (Heel and Schatz, 2017; van Heel and Schatz, 2005). Therefore, we used an independent method based on identification of features with sharp edges to verify our FSC resolution measurements (Mokso et al., 2007). This edge-fitting method produced resolution measurements consistent with those measured via FSC (Fig. S1d-f), suggesting that the FSC algorithm produces accurate measures of resolution that can be interpreted as the fidelity of sharp edges or small features in the image volumes.

### Correlative X-ray imaging with EM and genetic labeling

To verify that XNH images faithfully reproduce the ultrastructure of biological samples, we collected thin-sections of samples after XNH imaging and imaged the same regions at higher lateral resolution with EM. We found that corresponding EM and XNH image data from the *Drosophila* leg nerve (50 nm pixels, Fig. 2a, Supplementary Video 3) and mouse cortex (100 nm pixels, Fig. 2b) have very similar characteristics, with membranes and mitochondria appearing as the most prominent dark features. Most of the neurites in the EM image are also resolved in XNH, and only the smallest processes are not resolved with XNH. We also verified that fine ultrastructure features, such as membranes, synaptic vesicles and post-synaptic densities, are preserved in the EM images after XNH imaging (Fig. 2b, inset). This demonstrates that our XNH sample preparation and imaging approach are compatible with post-hoc, serial section EM, so that an XNH imaged volume can be imaged at higher resolution with EM, for identification of synapses or other fine ultrastructural features.

**Figure 2:**
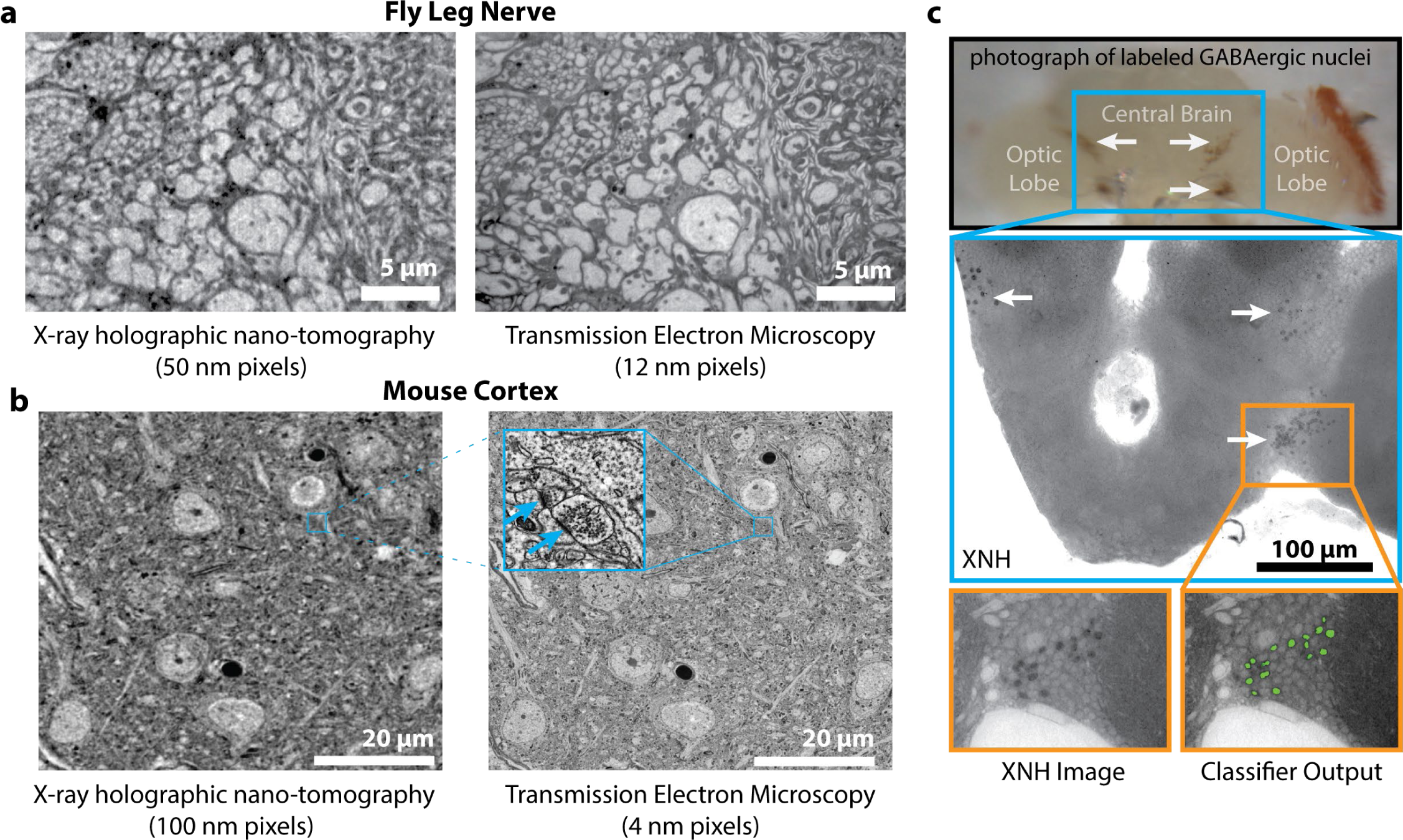
Ultrastructural Details and Genetic Labeling in X-ray Nano-Holography Images. **(a)** Comparison of corresponding X-ray (50 nm pixels) and TEM (12 nm pixels, 100 nm thick section) images taken from the *Drosophila* main leg nerve. Despite the difference in resolution, most of the motor and sensory axons are resolved in the X-ray image. **(b)** Comparison of corresponding X-ray (100 nm pixels) and TEM (4 nm pixels, 45 nm thick section) images taken from mouse somatosensory cortex. Inset: detail view of EM image showing chemical synapses (arrows) that are well-preserved after XNH imaging. **(c)** Top: photograph of fly brain with GABAergic nuclei labeled with APEX2 (arrows). Middle: XNH images (120 nm pixels, 15 µm thick minimum intensity projection) of the same fly brain, showing clusters of dark, APEX2 labeled GABAergic cell nuclei (arrows). Bottom: XNH virtual slice (120 nm thick) and output from an automated Random Forest image classifier trained to detect labeled cells (green).

Based on the similarity between XNH and EM images, we reasoned that cell type-specific labeling strategies previously developed for EM could be adapted for XNH. EM-visible labeling of specific cell types has been achieved using peroxidases that deposit dense precipitates inside the organelles of genetically-defined cell populations (Atasoy et al., 2014; Zhang et al., 2019). We developed a fly reporter line that targets the peroxidase APEX2 (Lam et al., 2015) to the nuclei of GABAergic neurons (see Methods). We show that APEX2 can successfully be detected in these flies with XNH imaging and that labeled cells can be automatically segmented using a Random Forest classifier with minimal training (Fig. 2c) (Sommer et al., 2011). These results demonstrate that the nuclear-APEX2 reporter line enables identification of genetically-defined cell populations within XNH images and suggest more generally that labeling strategies developed for EM can also be detected with XNH.

### Millimeter-scale XNH imaging of a *Drosophila* leg at single-neuron resolution

To image large regions with complex topology at high resolution, we stitched multiple, overlapping XNH scans to image the majority of a *Drosophila* leg (coxa, trochanter, femur, and tibia segments) and the associated first thoracic (T1) neuromere in the VNC (Fig. 3a,b). This dataset consisted of 12 individual scan volumes at pixel sizes between 50 and 200 nm, including a length of the main leg nerve totaling more than 1.4 mm (Fig. 3c,d, Supplementary Table 2, Supplementary Video 4). Because each XNH scan takes about 4 hours to acquire, this entire dataset was collected in less than 2 days. The 12 scans were aligned together to produce a continuous dataset encompassing the leg and the neural circuit in the VNC that controls this leg’s movements (Fig. 3c). Representative 2D slices (Fig. 3e-i) show that many axons in the main leg nerve can be clearly resolved and reconstructed. Because the surrounding musculature, exoskeleton, and sensory structures are also visible, individual neurons can be mapped from their peripheral sources and targets back to the VNC.

**Figure 3:**
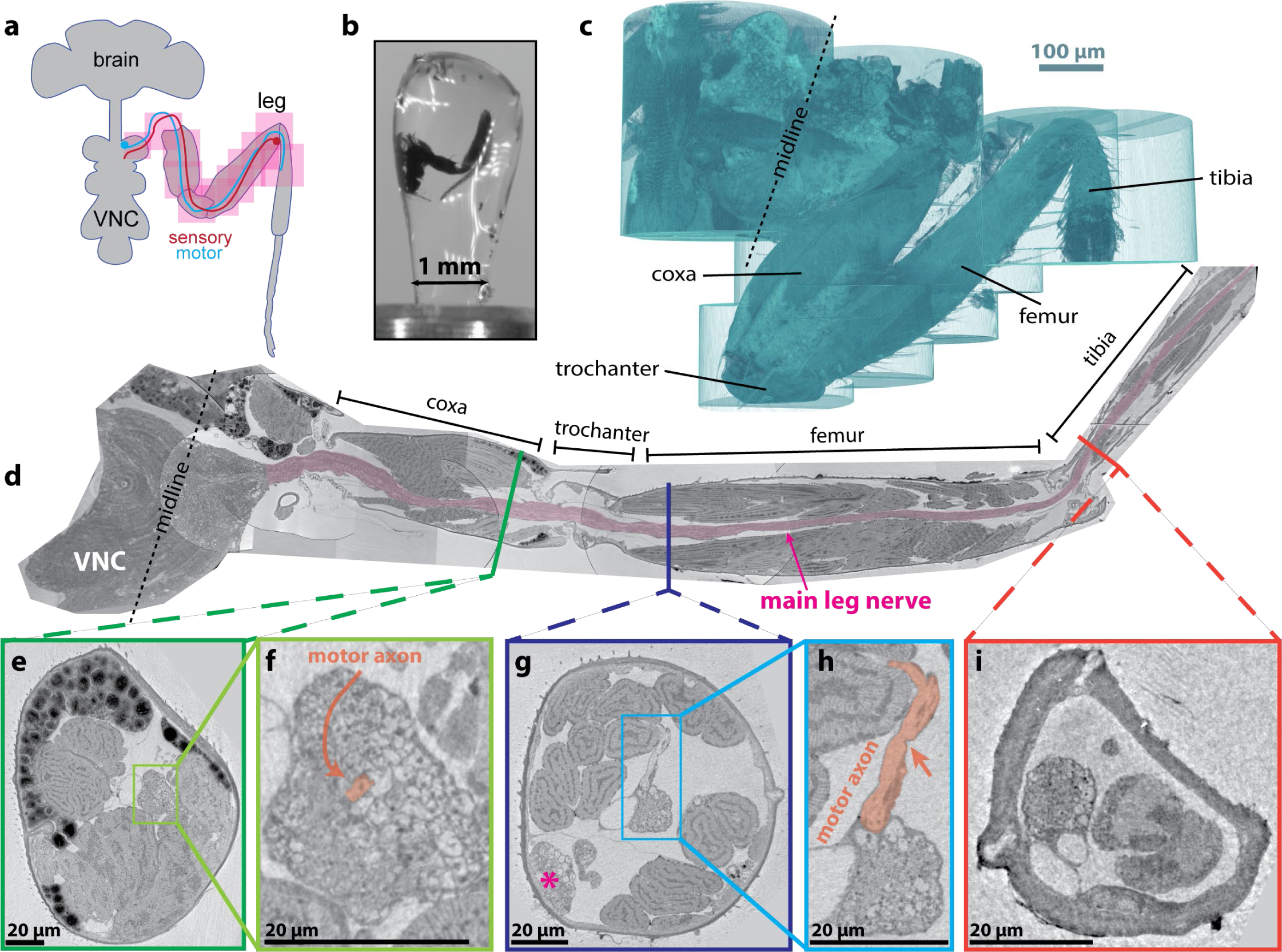
Millimeter-scale XNH imaging of a *Drosophila* leg at single-neuron resolution. **(a)** Schematic of XNH imaging strategy: 12 XNH scans were tiled along the front leg (Supplementary Table 2), including the T1 neuromere of the ventral nerve cord (VNC), the coxa, trochanter, femur, and tibia segments of the leg. Most scans were recorded at 75 nm pixel size, a resolution at which motor axons and large sensory axons can be traced through the leg. **(b)** Photograph of sample after heavy-metal staining, embedding, and mounting for XNH imaging. **(c)** 3D rendering of scan volume: individual scans were stitched together to form a contiguous volume of the leg (see Methods). **(d)** Computationally unfolded cross-section of the scan volume, following the main leg nerve over 1.4 mm from the VNC throughout the coxa, trochanter, femur, and tibia segments. **(e)** Cross-section through the coxa. Fats, muscles, and neurons are clearly visible. **(f)** Detailed view of nerve within the coxa. The indicated motor neuron is the same neuron as in (h) **(g)** Cross-section through the femur. Asterisk denotes the femoral chordatonal organ, a proprioceptive sensory structure (Mamiya et al., 2018). **(h)** Detailed view of nerve within the femur, including a motor axon branching off to innervate a muscle (arrow). **(i)** Cross-section through the tibia. Fewer large-diameter motor axons are visible in this cross-section compared to (f) and (h), as most motor axons have left the nerve to innervate muscles in more proximal leg segments.

### Reconstruction of Individual Neuron Morphologies

Until now, tracing of individual neuron morphologies from X-ray image data has been only possible through sparse labeling (Fonseca et al., 2018; Mizutani et al., 2013; Ng et al., 2016; Shahbazi et al., 2018). However, our results suggest that XNH image volumes contain sufficient signal to noise and spatial resolution to resolve and reconstruct dense populations of neurons without specific labeling. To test this, we reconstructed individual motor and sensory axons in high resolution scans of the *Drosophila* VNC (50 nm pixels, Supplementary Video 3) via manual skeletonization, a process where human annotators trace a wire-frame model of the neuronal processes (Saalfeld et al., 2009; Schneider-Mizell et al., 2016). A total of 108 neurons were seeded from their axons in the front leg nerve and were traced into the VNC (Fig. 4a-d). The reconstructed neurons included both motor neurons that send their axons into the leg to innervate muscles and sensory neurons that send signals from sensory organs in the leg to the VNC. We were able to trace neurons and capture the major branching patterns of their primary axons in the VNC. Based on these branching patterns, we were able to classify the neurons into several morphological clusters (Fig. 4e) and identify the dorsally-branching clusters as motor neurons (Baek and Mann, 2009; Brierley et al., 2012) and the ventrally-branching clusters as sensory neurons (Tsubouchi et al., 2017). In some cases, we were also able to associate the neurons with specific sensory organs, such as the campaniform sensilla and femoral chordatonal organs (Mamiya et al., 2018). We observed that axons are spatially organized in the nerve, such that axons from neurons from the same morphological cluster tend to also physically cluster together within the nerve (Fig. 4c, inset). These results demonstrate that XNH enables morphological classification of cell types, providing insight into the organizational principles of circuits in the central and peripheral nervous systems.

**Figure 4:**
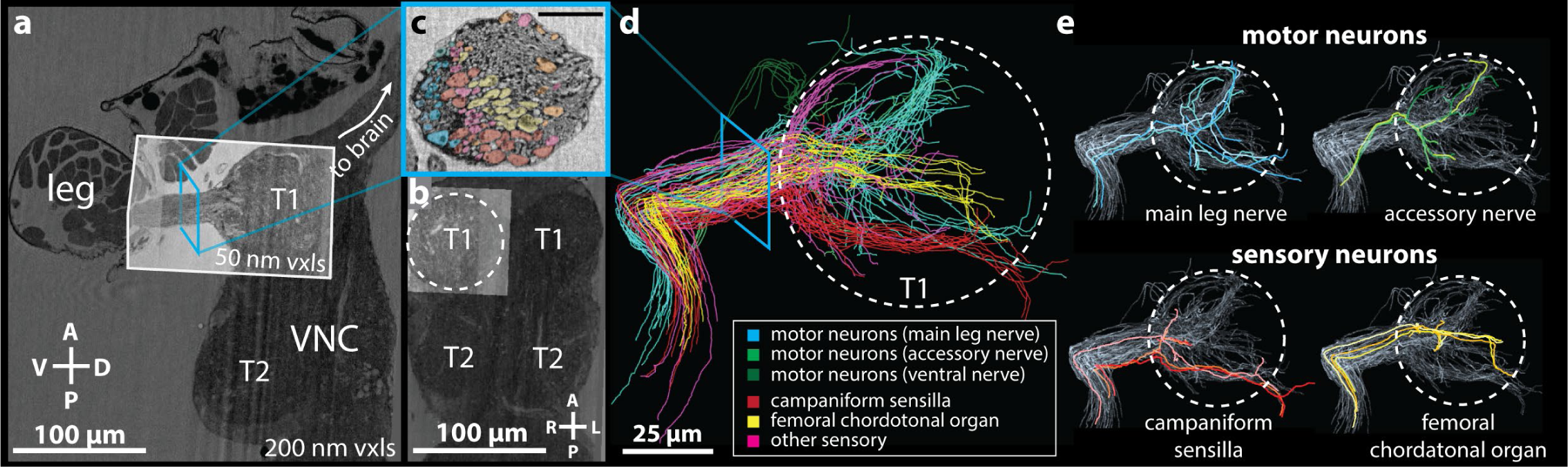
Reconstruction of Individual Neuron Morphologies. **(a-b)** Overview of XNH image volume encompassing the anterior half of the VNC and the first segment of a front leg of an adult *Drosophila* (200 nm voxels). A smaller, high resolution volume (50 nm voxels, highlighted) centered on the first thoracic (T1) neuromere of the VNC and including the initial segment of the leg nerve was used for tracing. **(c)** Virtual slice though the main leg nerve at the location indicated by the cyan square in (a) and (d). Colors of the individual neuron cross-sections correspond to the neuron type classifications shown in (d) Scale bar: 10 µm. **(d)** Left: 108 individual axons were manually skeletonized and clustered based on their morphologies. **(e)** Example neuron morphologies for motor (top) and sensory (bottom) neuron clusters. Axons arborizing dorsally in the VNC were identified as motor neurons (Baek and Mann, 2009; Brierley et al., 2012) and those arborizing ventrally as sensory neurons (Tsubouchi et al., 2017). Two major subtypes of leg sensory neurons, campaniform sensilla and chordotonal neurons, were identified based on arborization pattern in the VNC (Mamiya et al., 2018).

### Automated Segmentation of Neuronal Morphologies using Convolutional Neural Networks

While manual tracing can produce targeted, high-quality reconstructions, it can be time-consuming. To accelerate neuronal reconstructions, we adapted an automated segmentation pipeline developed for EM (Funke et al., 2017), and applied it to the XNH image data (see Methods). The goal of automated segmentation is to assign voxels in the image volume to neurons. Unlike skeletonization (Fig. 4), segmentation allows the full geometry of the neurons to be reconstructed in 3D. Broadly speaking, the pipeline consists of two major steps: affinity prediction and agglomeration. In the affinity prediction step, a CNN (Fig. 5a) is used to convert the image data into an affinity graph, which quantifies locally how likely adjacent voxels are to be part of the same object (Fig. 5b,c). During the subsequent agglomeration step, voxels are first grouped into supervoxels using a watershed algorithm, which are then progressively merged into larger objects based on the predicted affinities along the contact area of adjacent supervoxels. The merging procedure is stopped at a user-defined threshold, giving rise to the final segmentation (Fig. 5d).

**Figure 5:**
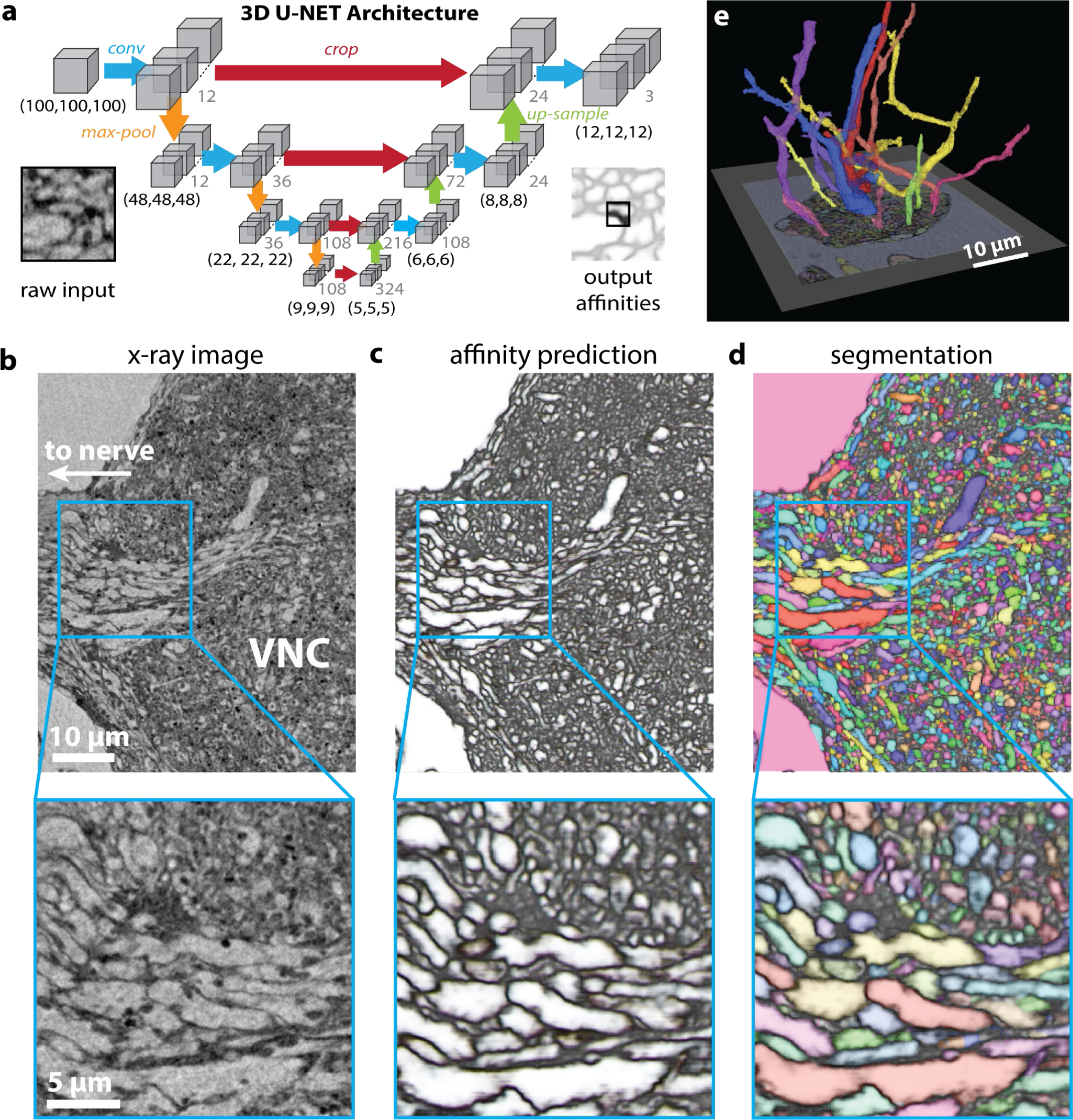
Automated Segmentation of Neuronal Morphologies using Convolutional Neural Networks (CNNs). **(a)** Schematic of U-NET CNN architecture used for automated segmentation (adapted from (Funke et al., 2017)). **(b-d)** Automated segmentation of XNH image volumes. **(b)** Raw XNH image data, recorded from *Drosophila* VNC. **(c)** Predicted affinities output by 3D U-NET corresponding to data shown in (b). For each voxel, the CNN calculates an affinity vector that quantifies how likely the pixel is grouped in the same neuron as neighboring pixels in z, y and x directions (plotted as RGB color components, respectively). In isotropic XNH data, affinities in different cardinal directions are usually similar, leading to images that appear mostly grayscale. These dark (low affinity) voxels are the basis for membrane predictions. **(d)** Segmentation of volume corresponding to data shown in (a) and affinities shown in (b). Each neuron is agglomerated into a 3D morphology based on the affinities. In this visualization, each neuron is colored with a unique color. **(e)** 3D visualization of automatically segmented neurons in the *Drosophila* VNC.

To assess whether automated segmentation can be used with XNH data to reconstruct neuronal morphologies, we applied automated segmentation to an XNH image volume encompassing the majority of a T1 neuromere and part of the main front leg nerve (Fig. 5e, Supplementary Video 5). Generally, the morphologies of the segmented neurons match the corresponding manual reconstructions, demonstrating that automated segmentation can generate dense single neuron morphologies from XNH data. However, in some instances, different voxels from the same neuron are erroneously labeled with different neuron IDs (split error) or two different neurons are erroneously combined into a single neuron ID (merge error). Such errors are also present in segmentations based on EM data, and are usually corrected via human proofreading. To determine how accurate the automated segmentation is, we compared it to manual (voxel-wise) segmentation for small test volumes and quantified split and merge errors using clustering similarity metrics (Fig. S2a, Supplementary Table 3). Comparing these error scores with published metrics from EM segmentation studies suggests that this automated segmentation workflow works similarly well with XNH data as with EM data and can be used to rapidly and accurately segment neurons.

## Discussion

### Advances in XNH imaging of neural circuits

Here, we present non-destructive, X-ray phase-contrast imaging of neural tissue with sufficiently high resolution and FOV to densely reconstruct individual neuronal morphologies. We have obtained these results by combining several innovative elements. The instrument developed at the ID16A beamline at the European Synchrotron (da Silva et al., 2017) can produce a highly brilliant and coherent X-ray probe focused to a spot below 30 nm, which we have used for holographic imaging. Another critical aspect is the accuracy of the rotational and translational stage movements to match the length scale of the expected resolution (< 100 nm); for this we employed active capacitive compensation to stabilize the sample (Villar et al., 2018). The sample itself must also remain stable throughout the measurements - a particular concern is heating and warping of the sample due to radiation absorption. We found that data quality depends on delivering an appropriate X-ray dose that is sufficient to produce clear images, but that does not warp the sample (Du and Jacobsen, 2018). Imaging in cryogenic conditions was a key development that mitigated sample warping and allowed us to reliably record data at sub-100 nm voxel sizes.

Sample properties are also important for image quality. Here, we chose to use heavy-metal staining protocols for EM, which enhance contrast in cell membranes, facilitate dense reconstruction of neurons, and allow the same samples to be imaged post-hoc with EM (Fig. 2a,b). However, with phase-contrast imaging, even unstained tissue can generate sufficient contrast to densely trace individual neurons (Fig. S1g, Supplementary Video 6). Phase-contrast imaging enables data acquisition from thick samples (> 1 mm), but image quality improves the closer the sample thickness is to the FOV. We found that a 300 μm thick sample of mouse cortex produces high-quality results (Fig. 1g), while a ~1 mm thick sample could still be imaged, but with a noticeable loss in resolution (Fig. 2b). Lastly, we developed a simple re-embedding method (Figs. 1a & 3b) to avoid rough edges on the surface of the sample, minimizing artifacts in the phase reconstructions.

### Neuron segmentation from XNH data

We demonstrate that XNH enables individual neuronal morphologies to be reconstructed in their dense circuit context. Here, we adapted reconstruction approaches developed originally for large-scale EM. Because XNH images appear qualitatively similar to EM images (Fig. 2), adapting these techniques was relatively straightforward. While XNH imaging currently cannot resolve the smallest neuronal branches (< 50 nm), reconstruction of a neuron’s large-diameter processes often clearly indicates its cell type (Schneider-Mizell et al., 2016). Indeed, we were able to identify pyramidal neurons in mouse cortex (Fig. 1g) and to differentiate sensory and motor neurons in the fly VNC (Fig. 4e).

The larger voxel size of XNH data relative to EM data offers several practical advantages. Larger voxels produce substantially smaller image files for a given tissue volume, which can be analyzed without specialized hardware infrastructure necessary for large-scale EM datasets. Furthermore, manual and automated analysis can be completed more rapidly, considering the reduced complexity of the task. Indeed, we deployed CNNs for automated segmentation in XNH volumes on timescales of a few hours (~ s/μm^3^ XNH compared with ~ 10s/μm^3^ for ssTEM).

### Outlook

XNH is complementary to and compatible with both LM and EM imaging. XNH offers superior resolution with denser labeling compared to most LM techniques, and can image samples too thick for LM without tissue clearing. XNH achieves lower resolution than volumetric EM, however, offers advantages in speed, convenience, and data handling. Since physical thin-sectioning is not necessary, XNH can image specimens that are difficult to cut reliably (such as a whole *Drosophila* leg). XNH image data also requires no section-to-section alignment, eliminating a step that continues to be a computational challenge for large-scale EM datasets.

By filling a gap between the resolutions of LM and EM, XNH can enable multiscale characterization of neural circuits that depend on both long-range connections and local computations, such as the *Drosophila* motor system and mammalian cortex. For example, a population of cortical neurons could be functionally imaged with LM, their long-range inputs mapped via XNH, and their local connectivity reconstructed using targeted EM.

At the resolutions achieved here, XNH is well-suited for mapping projectomes (i.e. atlases of all large-caliber connections between brain regions). Previously, single-neuron-resolution projectomes have been obtained using large-scale EM or built up from sparse fluorescent labeling (Chiang et al., 2011; Hildebrand et al., 2017); however, XNH represents a less laborious approach. A few XNH scans can encompass the whole brain of small model organisms such as adult *Drosophila* (Fig. 1), and in a typical beamline experiment (1-2 weeks), the entire brain of the larval zebrafish or larval tadpole could be mapped. With technical upgrades to increase imaging throughput, producing projectomes of small mammalian brains or a mouse brain may be possible within the next few years (Mikula, 2016).

It is worth noting that the results demonstrated here are still far from any theoretical resolution limits for hard X-rays, which have wavelengths on the angstrom scale. In practice, XNH resolution is limited by focusing optics, mechanical stability and precision of stage movements, sample warping and performance of reconstruction algorithms, rather than by fundamental physical limits. The upgrade of the European Synchrotron (to be completed in 2020), along with planned improvements to X-ray optics and detectors, will likely improve imaging resolution and throughput. Future technical advances may allow XNH to resolve the thinnest neuron branches and the synapses between them, opening a wide array of applications in mapping neuronal circuit connectivity.

## Acknowledgements

The authors acknowledge Jan Funke for providing code and assistance with automated segmentation; Julio da Silva for providing code and assistance for Fourier Shell Correlation measurements; John Tuthill and Tony Azevedo for discussions and advice regarding *Drosophila* motor systems; Norbert Perrimon, Matt Pecot, and Haluk Lacin for providing fly lines; Rick Fetter and Andrew Thompson for sample preparation advice; Rachel Wilson and Hannah Somhegyi for discussion and advice; Thedita Pedersen for preprocessing and alignment of image data; Lia DeCoursey, Rholee Xu, and Thedita Pedersen for neuron tracing; and Jimin Shin, Wei-Wei Lou, Julie Han, Mingguan Liu, Yumin Hu, and Rholee Xu for manual annotation of ground truth segmentation for CNN training.

## Data and materials availability

Image data and custom code will be made publicly available upon publication of the manuscript. In the interim, to request access to the data or custom code, please contact.

