## Supplementary Material for "Dense neuronal reconstruction through X-ray holographic nano-tomography"

This document includes:

1. Supplementary Tables
2. Descriptions of Supplementary Videos
3. Supplementary Figures
4. Materials and Methods

### Supplementary Tables

| Sample | Pixel Size | Scan Size | Scan Size (Ext FOV) | Measured Resolution | Measured Resolution |
|---|---|---|---|---|---|
| | (nm) | (µm) | (µm) | median [inter-quartile range] (nm) | voxels |
| mouse cortex | 30 | 61 | 96 | 87 [-4, +5] | 2.9 [-0.1, +0.2] |
| mouse cortex | 40 | 82 | 129 | 131 [-5, +6] | 3.3 [-0.1, +0.2] |
| mouse cortex | 40 | 82 | 129 | 134 [-3, +5] | 3.3 [-0.1, +0.1] |
| *Drosophila* VNC | 50 | 102 | 161 | 139 [-5, +5] | 2.8 [-0.1, +0.1] |
| *Drosophila* brain | 50 | 102 | 161 | 174 [-11, +10] | 3.5 [-0.2, +0.2] |
| *Drosophila* leg | 75 | 154 | 241 | 198 [-8, +10] | 2.6 [-0.1, +0.1] |
| *Drosophila* brain | 100 | 205 | 322 | 222 [-9, +10] | 2.2 [-0.1, +0.1] |
| *Drosophila* brain | 120 | 246 | 386 | 183 [-8, +16] | 1.5 [-0.1, +0.1] |

**Supplementary Table 1**: List of XNH scans included in resolution quantification (Figs. 1c, S1).

| Scan number | Contents | Pixel Size |
|---|---|---|
| | | (nm) |
| 1 | overview of VNC T1s | 160 |
| 2 | VNC, left T1 | 50 |
| 3 | overview centered on VNC left T1 nerve and top of coxa | 160 |
| 4 | left T1 nerve and top of coxa | 75 |
| 5 | Middle of coxa | 75 |
| 6 | Bottom of coxa and whole trochanter | 75 |
| 7 | Bottom of femur | 75 |
| 8 | Lower-middle of femur | 75 |
| 9 | Middle of femur | 75 |
| 10 | Upper-middle of femur | 75 |
| 11 | Femur-tibia joint | 75 |
| 12 | Upper 2/3rds of tibia | 100 |

**Supplementary Table 2**: List of XNH scans included in *Drosophila* leg dataset (Fig. 4).

| Dataset | Voxel Size (nm) | Method | VOI sum |
|---|---|---|---|
| XNH fly VNC test vol 1 (this work) | (50,50,50) | U-NET MALA | 1.231 |
| XNH fly VNC test vol 2 (this work) | (50,50,50) | U-NET MALA | 1.470 |
| FIBSEM (FIB-25) | (8,8,8) | U-NET MALA | 1.071 |
| | | U-NET | 1.524 |
| | | FlyEM | 1.952 |
| | | CELIS | 1.634 |
| | | CELIS+MC | 1.266 |
| ssTEM (CREMI) | (4,4,40) | U-NET MALA | 0.606 |
| | | U-NET | 1.524 |
| | | LMC | 0.868 |
| | | CRunet | 1.470 |
| | | LFC | 1.225 |

**Supplementary Table 3**: Quantification of segmentation error (VOI sum). Comparison values for EM datasets from (Funke et al., 2017). **Note:** a precise quantitative comparison of segmentation performance on EM and XNH data is difficult because human-annotated, ground truth segmentation of XNH data excludes areas where features are too small to resolve. Thus, cluster-based error metrics (such as VOI index as used in Fig. S2) may not be directly comparable to what has been reported for EM. Nevertheless, it is clear that automated segmentation algorithms can be successfully applied to XNH data.

#### Descriptions of Supplementary Videos

**Supplementary Video 1:** An XNH scan of an adult *Drosophila* brain at a voxel size of 120 nm and a measured resolution of 183 nm. At this resolution, many individual neurons can be tracked as they travel between different brain regions. The field of view encompasses the entire central brain and part of both optic lobes, allowing brain-wide projections to be mapped.

**Supplementary Video 2:** An XNH scan of mouse somatosensory cortex (layer 5) at a voxel size of 30 nm and a measured resolution of 87 nm. At this resolution, all myelinated axons and most unmyelinated axons and dendrites are resolved. Many subcellular features, most notably mitochondria and endoplasmic reticulum, are resolved.

**Supplementary Video 3:** An XNH scan of the adult *Drosophila* VNC, encompassing the T1 neuromere that controls movements of a front leg. This single scan at 50 nm voxels captures the majority of the T1 neuromere with sufficient resolution (139 nm) to reconstruct single neurons and identify different types of sensory and motor neurons (see Fig. 4). Body wall muscles surrounding the neuromere, as well as the motor neurons innervating those muscles, are also visible.

**Supplementary Video 4:** An XNH scan of part of the adult *Drosophila* leg at a voxel size of 75 nm and a measured resolution of 198 nm. The inset shows a zoom-in on the leg nerve, which contains motor neurons (large diameter axons) traveling out to muscles, as well as sensory neurons (variable diameter depending on type, but the majority are too small to resolve) bringing mechanosensory information to the central nervous system. The structural arrangement of muscles, fats, and joints are visible (see Fig. 3).

**Supplementary Video 5:** Automated segmentation of a portion of the adult *Drosophila* VNC (same data as Video 3 and Fig. 4). Left, each neuron is labeled by a distinct color, though some errors remain where different neurons share the same color, or two branches are colored differently despite belonging to the same neuron. Right, volumetric view of selected large-diameter neurons.

**Supplementary Video 6:** An XNH scan at 105 nm voxel size of an adult *Drosophila* brain prepared without any heavy metal staining. Phase-contrast imaging enables this type of soft tissue to be imaged with reasonable contrast. As with stained samples (Video 1), many individual neurons can be reconstructed traveling within and between brain regions.

### Supplementary Figures

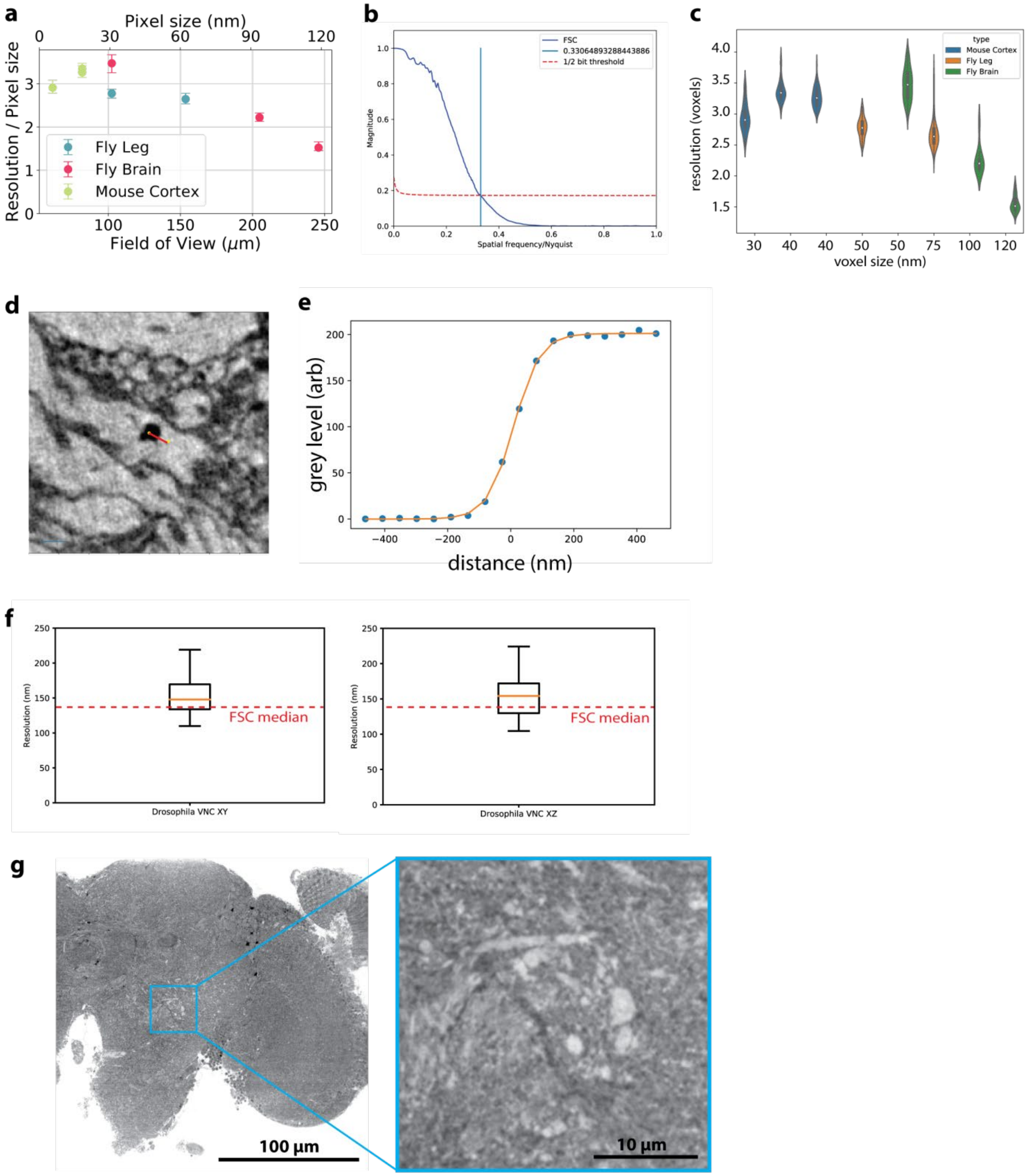


**Figure S1: (a)** Quantification of resolution of XNH scans measured using Fourier Shell Correlation (FSC), plotted in units of pixels. As the pixel sizes increase, the resolution per pixel improves. **(b)** Representative FSC curves shown with the half-bit threshold. The intersection between the FSC curve and the threshold is the measured resolution. **(c)** Distribution of resolution measurements for voxel subvolumes (~$250^3$ chunks, see Methods) shown for XNH scans shown in Fig. 1h. **(d-e)** Example line scan used for edge-fitting approach for measuring resolution (see Methods). The measured resolution is parameterized from a best-fit to a sigmoid function. **(f)** Distribution of edge-fitting resolution measurements for an XNH scan taken in the *Drosophila* VNC (50 nm voxels) for XY and XZ planes. The median resolution measured via FSC is shown for comparison. **(g)** XNH data (105 nm vxls) of a Drosophila brain that did not undergo heavy metal staining. Even in unstained soft tissue, phase-contrast imaging provides enough signal that single neurons can still be resolved. FOV encompasses the optic lobe and half of the central brain. See Supplementary Video 5 and Methods.

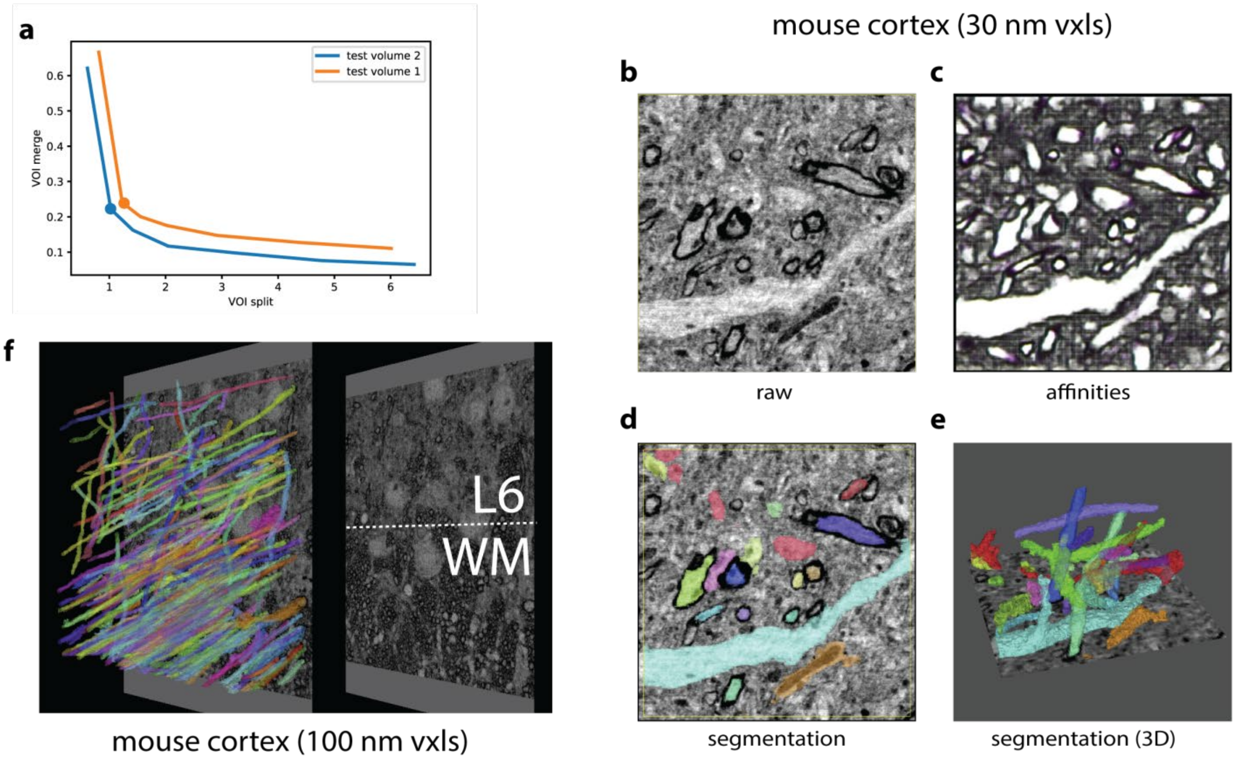


**Figure S2: Automated Segmentation of Neuronal Morphologies using Convolutional Neural Networks. (a)** Variation of information error quantification of segmentation based on XNH Drosophila VNC data (50 nm pixels). The agglomeration threshold θ was used to parameterize the curve. The point of minimal sum of VOI, corresponding in these data to θ = 0.7, are indicated by the markers in the plot. A comparison to the VOI sum metric in other datasets is shown in Supplementary Table. 3. **(b-e)** Automated segmentation of XNH data in mouse cortex (primary somatosensory, layer 5, 30 nm voxels). **(b)** Raw data **(c)** Affinities (zyx corresponding to RGB colors). **(d)** selected segmentation labels corresponding to (e). **(e)** Selected 3D renderings of segmented neuron fragments. **(f)** Large FOV segmentation of myelinated axons in the white matter below mouse parietal cortex. Segmentation of such myelinated axons allows tracing of long-range inputs between brain areas at single-cell resolution.

### Materials and Methods

#### Sample preparation

Tissue samples were prepared for XNH imaging using protocols for electron microscopy (EM), including fixation, heavy metal staining, dehydration, and resin embedding (Hua et al., 2015; Zheng et al., 2018). For heavy metal staining, we used an enhanced rOTO protocol (Hua et al., 2015) for all mouse samples and some fly samples. For other fly samples, we modified the protocol to increase (Zheng et al., 2018) or decrease heavy metal staining, but these variations did not have a large effect on XNH image quality. Following staining, samples were dehydrated in a graded ethanol series and embedded in either TAAB Epon 812 (Canemco) or LX112 (Ladd Research Industries) resin. Resin-embedded samples were polymerized at 60ºC for 2-4 days. Polymerized samples were trimmed down to a narrow (1 to 2 mm diameter) rod using either an ultramicrotome or a fine saw, then glued to an aluminum pin. To smooth rough surfaces on the samples, which introduce noise into the XNH images, mounted samples were covered in a small droplet of resin, and the droplet was polymerized at 60ºC for 2-4 days.

For labeling APEX2-expressing cells (see below), we performed 3,3'-diaminobenzidine (DAB) staining after fixation and before heavy metal staining. Briefly, the nervous system from an adult female was dissected, fixed (2% glutaraldehyde, 2% formaldehyde solution in 100mM sodium cacodylate buffer with 0.04% CaCl2) at room temperature for 75 minutes then moved to 4ºC for overnight fixation in the same solution. The following day, the sample was washed in cacodylate buffer, then 50mM glycine in cacodylate buffer, then cacodylate buffer again. To stain the APEX2-expressing nuclei, the sample was incubated with 0.03% DAB in cacodylate buffer for 30 minutes, then $H_2O_2$ was added directly to the incubating samples to reach an $H_2O_2$ concentration of 0.003% (Zhang et al., 2019). The reaction was allowed to continue for 30 minutes, after which the sample was washed in cacodylate buffer and inspected for visible staining product. Clusters of small brown puncta corresponding to the labeled nuclei were faintly visible (Fig. 2c, top panel). The DAB and $H_2O_2$ incubations were repeated once more to increase the staining intensity. The sample was subsequently stained with 3-amino-1,2,4-triazole-reduced osmium (Zheng et al., 2018) and uranyl acetate, then dehydrated and embedded in LX112 resin.

The unstained sample (Fig. S1g, Supplementary Video 6) was fixed in Karnovsky's fixative, dehydrated in ethanol, cleared with xylene, and embedded in paraffin. The embedded sample was trimmed down to a narrow (1 to 2 mm diameter) rod using a scalpel, and inserted into a hollow aluminum pin with a 0.8 mm inner diameter for imaging.

#### Experimental setup and data acquisition

XNH imaging was performed at beamline ID16A at the European Synchrotron in Grenoble, France. The end-station was placed 185 m from the undulator source for improved coherence. The X-ray beam was focused using fixed curvature, multilayer coated Kirkpatrick-Baez mirrors into a spot measuring about 15 nm at X-ray energy of 33.6 keV (da Silva et al., 2017) and 30 nm at 17 keV. The photon flux was on the order of 1-4 x $10^{11}$ ph/s.

The sample stage and X-ray focusing optics were placed in a vacuum chamber (pressure ~$10^{-8}$ mbar) and a liquid nitrogen based cryogenic system was integrated inside the stage, keeping the sample at 120 K during imaging. For cryogenic imaging, the samples were transferred into the vacuum chamber with a Leica cryo-shuttle. The samples were placed on a high-precision rotation stage (Villar et al., 2018) downstream of the beam focus, and intensity projections (i.e. holograms) were recorded using a FReLon 4096 x 4096 pixel CCD detector (Labiche et al., 2007) with 2x binning, lens-coupled to a 23 μm thick GGG:Eu scintillator.

Holograms were recorded at four different sample-to-focus distances and combined together to obtain a phase map through a phase retrieval algorithm (Cloetens et al., 1996; Mokso et al., 2007). In practice, four tomographic scans were recorded sequentially at different distances by rotating the sample around a vertical rotation axis. To eliminate ring artifacts in tomographic reconstructions, the samples were laterally displaced at each rotation angle by a randomly-determined distance of up to 25 pixels using high-precision piezoelectric actuators (Hubert et al., 2018). During data processing, corresponding sets of four holograms were aligned and brought to the same magnification before applying a phase retrieval algorithm. The divergent beam gives geometrical magnification $M = (z_1 + z_2)/z_1$ with $z_1$ = focus-to-sample distance and $z_2$ = sample-to-detector distance. Therefore, the pixel size and the corresponding FOV are proportional to $z_1$ when the detector position is fixed (Fig. 1a).

For each tomographic scan of the mouse cortex, 1800 projections were recorded with exposure times of 0.1 s at X-ray energy 17 keV and 0.35 s at 33.6 keV. For the *Drosophila* scans at 17 keV, 2000 projections were collected with 0.2 s exposure times.

#### Image reconstruction

The recorded holograms were initially preprocessed to compensate for distortions and noise specific to the optics and detector, and normalized with the empty beam (Hubert et al., 2018). For each rotation angle, the four holograms corresponding to different propagation distances were aligned and brought to the same magnification. Normally, the holograms are cropped to the smallest FOV, corresponding to the targeted pixel size. In order to obtain an extended FOV, the information from the three larger FOVs at lower resolutions was integrated in the reconstruction as well. From these sets of aligned holograms, phase maps were obtained through an iterative algorithm. The initial approximations of the amplitude and phase were obtained through a method based on (Paganin et al., 2002), adapted for multiple propagation distance holograms (Yu et al., 2018). For regularization we used the ratio between the refractive index decrement $\delta$ and the absorption index $\beta$ corresponding to Osmium ($\delta/\beta = 27$ for X-ray energy 33.6 keV and $\delta/\beta = 9$ for 17 keV). At each iterative step, the amplitude term was kept constant and the phase term was updated. Typically, 10 iterations were sufficient for the phase term to converge. Computation time was approximately 15 minutes per phase map (single CPU node). Computation of the phase maps was done in parallel by treating the holograms for each rotation angle independently.

#### Resolution Measurements

##### Fourier Shell Correlation

One of the methods used to measure the effective resolution in the tomographic reconstructions was Fourier Shell Correlation (FSC) (Harauz and van Heel, 1986). This consists of splitting the data into two separate image volumes, assuming independent noise, and measuring the normalized cross-correlation coefficient between the two volumes over corresponding shells in Fourier space. This metric was measured across shells, and then the intersection between the FSC line and a horizontal threshold (van Heel and Schatz, 2005) was used to determine the resolution (Fig. S1b). For our measurements, we used the ½-bit threshold.

To ensure that the noise from each volume is independent (van Heel and Schatz, 2005), the two image volumes were generated independently from half of the phase maps (even and odd projections were separated). Once the volumes were reconstructed, FSC was applied in small chunks throughout the volume. The size of these chunks was selected by increasing the size until the FSC metric was stable (typically ~ $(250\ \text{vxls})^3$). This ensured that we do not select a chunk size that generates artificially high resolution values. Larger chunks (up to $(1000\ \text{vxls})^3$) remained stable but took longer to compute. Once the chunk size was selected, chunks were evenly spaced across the volume, ignoring parts of the volume with only empty resin. Once the resolutions for all chunks were measured, the median and IQR were determined (Fig. 1h, Fig. S1a,c).

#### Edge-Fitting

For measuring resolution by edge-fitting, line paths perpendicular to sharp edges in the image volumes were annotated manually using CATMAID (Fig. S1d) (Saalfeld et al., 2009; Schneider-Mizell et al., 2016). The image intensity along these line paths (either in xy or xz planes) were then calculated using the Pymaid python API (https://github.com/schlegelp/PyMaid). The points along the line paths were fit to the sigmoid function:

$$p_3 + \frac{p_0}{2}\left(1 - \tanh\left(\frac{x - p_1}{p_2}\right)\right)$$

Where $x$ is the length along the line path and $p_{1-4}$ are free parameters determined by nonlinear regression. (Fig. S1e).

Given a fit to an edge, the measured resolution is given by:

$$\text{res} = \text{asech}\left(\frac{2}{\sqrt{2}}|p_2|\right)$$

The fits to each line path were inspected and poor fits were refit with different initial parameters or removed. Edge-fitting was used in the 50 nm voxel dataset taken in the *Drosophila* VNC (see Supplementary Table 1). In total, 33 measurements were taken in XY planes and 20 measurements were taken in XZ planes (Fig. S1f,g)

#### Post-Hoc EM imaging

After completing XNH imaging, samples were re-embedded in a block of resin and trimmed for thin-sectioning. Serial thin sections (45-100 nm) were cut using a 35 degree diamond knife (Diatome) and collected onto LUXFilm-coated copper grids (Luxel Corp.). Sections were imaged on a JEOL 1200EX transmission electron microscope (80 kV accelerating potential, 1500x mag), and images were acquired with a 20 MPix camera system (AMT Corp.) at 4-12 nm pixels.

The XNH virtual slices shown in Fig. 2a,b were rotated and aligned to match the corresponding EM micrographs. For Fig. 2a, the XNH dataset of the fly VNC was rotated in Neuroglancer (https://github.com/google/neuroglancer) to match the orientation of the transmission EM (TEM) section. The TEM image of the leg nerve was elastically aligned to a single matching image taken from the XNH dataset using AlignTK (https://mmbios.pitt.edu/aligntk-home). For Fig. 2b, the XNH dataset of mouse cortex was aligned to the TEM micrograph via an affine transformation based on manually annotated correspondence points (annotated using BigWarp https://imagej.net/BigWarp).

#### Generation of Nuclear-APEX2 Flies

To target APEX2 to the nucleus, we fused a targeting sequence consisting of a methionine and 38 amino acids of the Stinger sequence (MSRHRRHRQRSRSRNRSRSRSSERKRRQRSRSRSSERRR) to APEX2. The targeting sequence was first cloned into the pENTR vector and subsequently cloned by recombination using the Gateway system into a destination vector (gift from Dr. Frederik Wirtz-Peitz) containing UAS-attR-sbAPEX2-3xMyc. The resulting UAS-NLS-APEX2-Myc construct was used to generate a transgenic line by direct injection using φC31 site-specific integration at the attP40 docking site on chromosome two.

#### Labeling of GABAergic nuclei with Nuclear-APEX2

Fly lines containing the transgenes Gad1-p65AD, UAS-CD8-GFP and elav-Gal4DBD, UAS-CD8-GFP (gifts from Dr. Haluk Lacin) were crossed with the nuclear-APEX2 fly described above to generate flies with genotype w; elav-Gal4DBD, UAS-CD8-GFP / UAS-NLS-APEX2-Myc; Gad1-p65AD, UAS-CD8-GFP / +. The nervous system from a 5-6 day old adult female was prepared for XNH imaging as described above.

#### Classifier for Detecting Labeled Cells

A 3D random forest pixel classifier was trained to detect APEX2-labeled cell bodies in a subset of the XNH dataset shown in Fig. 2c. The model was created, trained, and deployed using ilastik (https://www.ilastik.org/). For training, a sparse set of pixel labels was interactively annotated for background pixels and labeled cell body pixels.

#### Image Volume Stitching

For each pair of XNH scan volumes with overlapping FOVs, correspondence points identifying the same feature in each scan were annotated manually using the ImageJ plugin BigWarp (https://imagej.net/BigWarp, (Bogovic et al., 2016)). Translation-rotation-scaling matrices were calculated based on least-squares fitting of these correspondence points (~10-20 pairs per image volume) using custom MATLAB code, then applied to each image volume using the ImageJ plugin BigStitcher (https://imagej.net/BigStitcher). To avoid blurring from misalignments in regions where two scans overlap, image volumes were combined without blending in overlapping regions (custom Python code).

#### Neuron Tracing

Manual tracing of neurons (Fig. 4) was performed by a team of 2-4 annotators using CATMAID (Saalfeld et al., 2009; Schneider-Mizell et al., 2016). Neuron morphologies were visualized using the CATMAID 3D viewer and manually clustered.

#### Automated Segmentation

We used an automated segmentation workflow based on a segmentation pipeline for TEM data (Funke et al., 2017). The pipeline consists of two major steps: affinity prediction and agglomeration. In the affinity prediction step, a 3D U-Net convolutional neural network (CNN) was used to predict an affinity graph from the image data. The value of the affinity graph at any given pixel represents the probability that adjacent pixels (in the x, y, and z axes) are part of the same object. Pixels in membranes should have low affinity values, where pixels within neurons should have high affinity to the pixels surrounding them (including pixels contained in organelles or other subcellular structures). To expedite training a CNN on XNH data, we leveraged a CNN first trained on ground truth EM volumes from the CREMI challenge (https://cremi.org/), followed by training augmentation with corrected segmentation predicted on XNH data. We began by downsampling the CREMI ground truth data (4 nm x 4 nm x 40 nm) to match the voxel size of the XNH data (50 nm x 50 nm x 50 nm). Then, starting from a candidate segmentation generated by this CREMI-trained network on XNH data, we manually corrected small subvolumes using Armitage (Google) and ITK-Snap (http://www.itksnap.org) to produce (semi-sparse) ground truth XNH training volumes. After augmented training on the XNH ground truth, the CNN was deployed across the entire XNH volume. Following affinity prediction, we agglomerated by progressively grouping voxels into larger objects based on the affinities connecting them to neighboring voxels. The final segmentation depends on a chosen agglomeration threshold. As training improved, this threshold was increased, such that larger objects could be segmented without introducing erroneous merges.
